# The mitochondrial genome of *Palythoa tuberculosa* (Esper, 1805) contains a novel ORF with unknown function

**DOI:** 10.1101/2025.09.26.678703

**Authors:** Yuki Yoshioka, Megumi Kanai, Tomofumi Nagata, Noriyuki Satoh

**Author notes:** Corresponding author: Yuki Yoshioka.

## Abstract

Recent advances in sequencing technology have enabled rapid and cost-effective recovery of mitochondrial genomes (mt-genomes), which used in phylogenetic and evolutionary studies. Among cnidarians, the order Zoantharia possesses the largest mt-genomes on average (20,976 bp). Here, we present the complete mt-genome of *Palythoa tuberculosa* (Hexacorallia: Zoantharia: Sphenopidae), which was 23,044 bp in length—approximately 2,000 bp larger than the zoantharian average. Synteny analysis revealed that this increase resulted from a novel open reading frame (ORF) located between *trnW* and *cox2*. This ORF was also present in the closely related species *P. caribaeorum*, indicating its origin in their common ancestor. Importantly, this represents the first case of a novel ORF insertion in zoantharians, clearly distinct from previously reported ORF extensions. While mt-genome expansions are occasionally observed in cnidarians, both the insertion sites and inserted elements vary among lineages, highlighting the lineage-specific mechanisms driving mitochondrial genome evolution in this group.

## Introduction

Since the advent of next-generation sequencing, obtaining complete mt-genomes has become much faster and more cost-effective. Animal mt-genomes are typically circular and range 14.5 to 19.5 Kbp in length. In cnidarians, the mt-genome sizes range 14,320 to 23,997 bp (NCBI Nucleotide database, as of August 29, 2025), with the subphylum Anthozoa having the highest number of mt-genomes registered in public database within the phylum Cnidaria. The mt-genome sizes of anthozoans are relatively stable and show a low standard deviation (Feng et al., 2023). Within Anthozoa, the order Zoantharia, an early-diverging lineage of Hexacorallia (Quattrini et al., 2020), possesses comparatively larger mt-genomes, with an average size of 20,976 bp (Feng et al., 2023). Most of the size variation among animal mt-genomes is attributable to differences in the amount of noncoding DNA, which is often unevenly distributed among intergenic regions (Lavrov and Pett, 2016).

The genus *Palythoa* Lamouroux, 1816 belongs to the family Sphenopidae (class Hexacorallia: order Zoantharia) and is well known to produce palytoxin, one of the most potent natural toxins, with an LD50 of 25-450 ng/kg body weight (Wiles et al., 1974). Among *Palythoa* species, *Palythoa tuberculosa* (Esper, 1805) consistently produces palytoxin and its congener 42-hydroxy-palythoxins (Aratake et al., 2016), making it a promising model species for studying palytoxin biosynthesis. Although a complete mt-genome sequence of *P. tuberculosa* was reported by Fourreau et al. (2023), gene annotation for this genome has not been made publicly available. Notably, the complete mt-genome of *P. tuberculosa* is 23,044 bp in length and is approximately 2,000 bp larger than the average of other zoantharian members, but the underlying cause of this genome size expansion has not been studied. In cnidarians, mt-genome size increase has been attributed to tRNA gene duplication (Chen et al., 2008), an insetion of open reading frame (ORF) (Flot et al., 2007), and/or the insertion of non-coding regions (Brockman and McFadden, 2012; Yoshioka et al., 2025a). In this study, we report the complete mitochondria genome sequences of *Palythoa tuberculosa* with gene annotations, and we perform comparative synteny analyses with several *Palythoa* species to reveal the evolutionary mechanism underlying its genome size expansion.

## Materials and method

We collected *P. tuberculosa* at Mizugama (sample IDs 42 and 82) and Sunabe (sample IDs 7 and 12), west coast of Okinawa Island, Okinawa, Japan, which were preserved in 99.5% ethanol until use. Species was identified based on the description by Reimer et al. (2014). To ensure species identification and sequence accuracy, we collected four individuals of *P. tuberculosa*. We extracted genomic DNA from preserved specimens using a Maxwell RSC Blood DNA Kit (Promega). Sequence libraries were constructed with a KAPA Hyper Prep Kit (KAPA Biosystems) according to the manufacturer’s protocol and sequenced on a NovaSeq 6000, with 250-bp paired-ends. Illumina sequence adaptors and low-quality sequences (quality cutoff=20) were trimmed with CUTADAPT v4.3 (Martin, 2011). Cleaned reads were assembled with GetOrganelle v1.7.7.0 (Jin et al., 2020). Mitochondrial gene annotation was performed with MITOS2 (Bernt et al., 2013), followed by manual curation. Complete mt-genomes with gene annotation were visualized with OrganelleGenomeDRAW (Greiner et al., 2019). Gene order was visualized with Clinker (Gilchrist and Chooi, 2021). Raw genomic DNA sequences used in this study were deposited in the DDBJ Sequence Read Archive under accession numbers DRR727336–DRR727339 (BioProject ID: PRJDB17204). Mitogenomes are available under accession numbers LC891139–LC891142.

We downloaded mt-genomes of all zoantharians used in this study from NCBI on 24 September 2024. Starting positions of whole-mt-genomes were adjusted across taxa and aligned using MAFFT v7.520 (Katoh et al., 2002) with the option (--auto). Poorly aligned regions in alignments were trimmed with ClipKit v2.2.4 (Steenwyk et al., 2020) with the option (-m gappy). Molecular phylogenetic analyses were carried out using IQ-TREE v2.2.2.6 (Minh et al., 2020) with the option (-m MFP; -B 1000). In the analysis, ModelFinder (Kalyaanamoorthy et al., 2017) (as packaged in IQ-TREE) was used to identify the best-fit substitution model. tRNAs were visualized with RNAfold web server (Gruber et al., 2008). Relative synonymous codon usage (RSCU) was estimated using RSCUcaller v0.99.0 (Maździarz et al., 2025).

Selective constraints on the protein coding genes (PCGs) in the mt-genome of *P. tuberculosa* were investigated. Ka (number of nonsynonymous substitutions per non-synonymous site), Ks (number of synonymous substitutions per synonymous site), and Ka/Ks were estimated for each gene using KaKs_calculator 2.0 (Wang et al. 2010). Values were estimated using a pairwise comparison between *P. tuberculosa* and the congeneric *P. mutuki* (GenBank accession number NC_046404). The Nei-Gojobori model was used during calculations to account for variable mutation rates across sequence sites.

## Results and Discussion

We obtained complete mt-genome sequences from four *P. tuberculosa* individuals, each with an assembled size of 23,044 bp (Figure 1). Sequence comparison among *P. tuberculosa* individuals revealed only a single nucleotide substitution in the intergametic region between *cob* and *cox3*, indicating high sequencing accuracy. The mt-genomes contained 13 PCGs (*atp6, atp8, cob, cox1*–*3, nad1*–*6*, and *nad4l*), two rRNA genes, and two tRNA genes (Figure 1), the composition of which was consistent with the previously reported one (Fourreau et al., 2023). All mitochondrial PCGs started with the start codon ATG as with other *Palythoa* species (Fourreau et al., 2023; Yoshioka et al., 2024). Genes *atp6, cox1*–*3, nad1*, and *nad2* terminated with stop codon ATG, while other genes terminated with TAA (Table 1). Notably, we discovered a novel ORF located between *trnW* and *cox2* in the mito-genome of *P. teberculosa*. This ORF encoded 665 amino acids, initiated with the codon ATG and terminated with TAA (Figure 1 and Supplementary Figure S1). Both tRNAs exhibited typical “cloverleaf” secondary structure, although *trnW* showed relatively shorter anticodon arm compared to *trnM* (Supplementary Figure S2). Longer intergenic regions, either between *cox2* and *nad4* (424 bp), between *cob* and *cox3* (265 bp), or both, may function as control regions (Supplementary Figure S3).

**Table 1.**
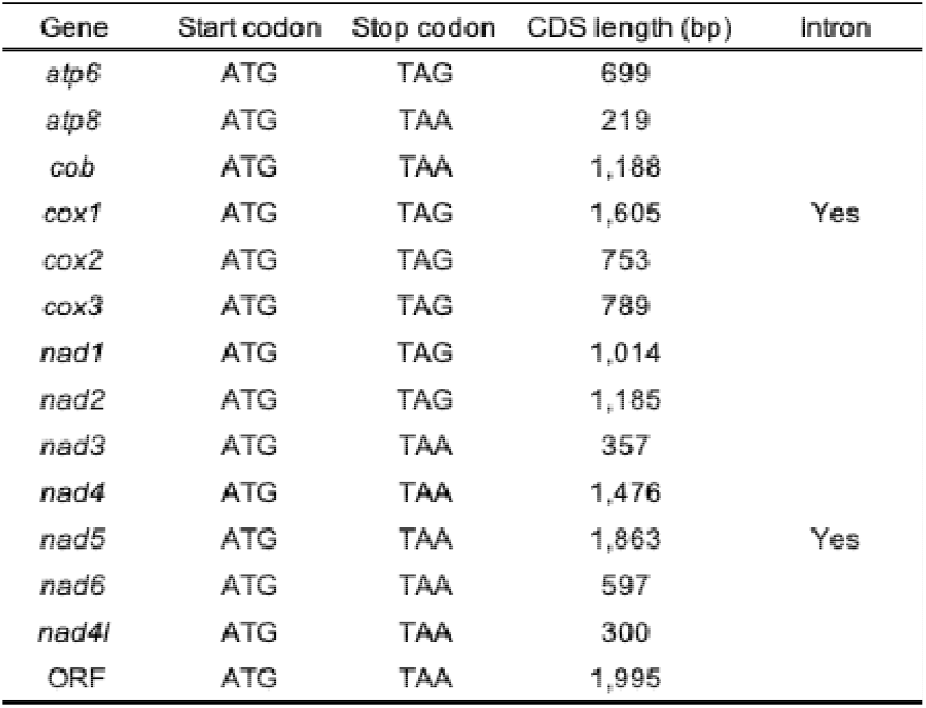
Start and stop codons and gene length of mitochondrial protein coding genes of *Palythoa tuberculosa*.

| Gene | Start codon | Stop codon | CDS length (bp) | Intron |
| --- | --- | --- | --- | --- |
| *atp6* | ATG | TAG | 699 |  |
| *atp8* | ATG | TAA | 219 |  |
| *cob* | ATG | TAA | 1,188 |  |
| *cox1* | ATG | TAG | 1,605 | Yes |
| *cox2* | ATG | TAG | 753 |  |
| *cox3* | ATG | TAG | 789 |  |
| *nad1* | ATG | TAG | 1,014 |  |
| *nad2* | ATG | TAG | 1,185 |  |
| *nad3* | ATG | TAA | 357 |  |
| *nad4* | ATG | TAA | 1,476 |  |
| *nad5* | ATG | TAA | 1,863 | Yes |
| *nad6* | ATG | TAA | 597 |  |
| *nad4l* | ATG | TAA | 300 |  |
| ORF | ATG | TAA | 1,995 |  |

**Figure 1.**
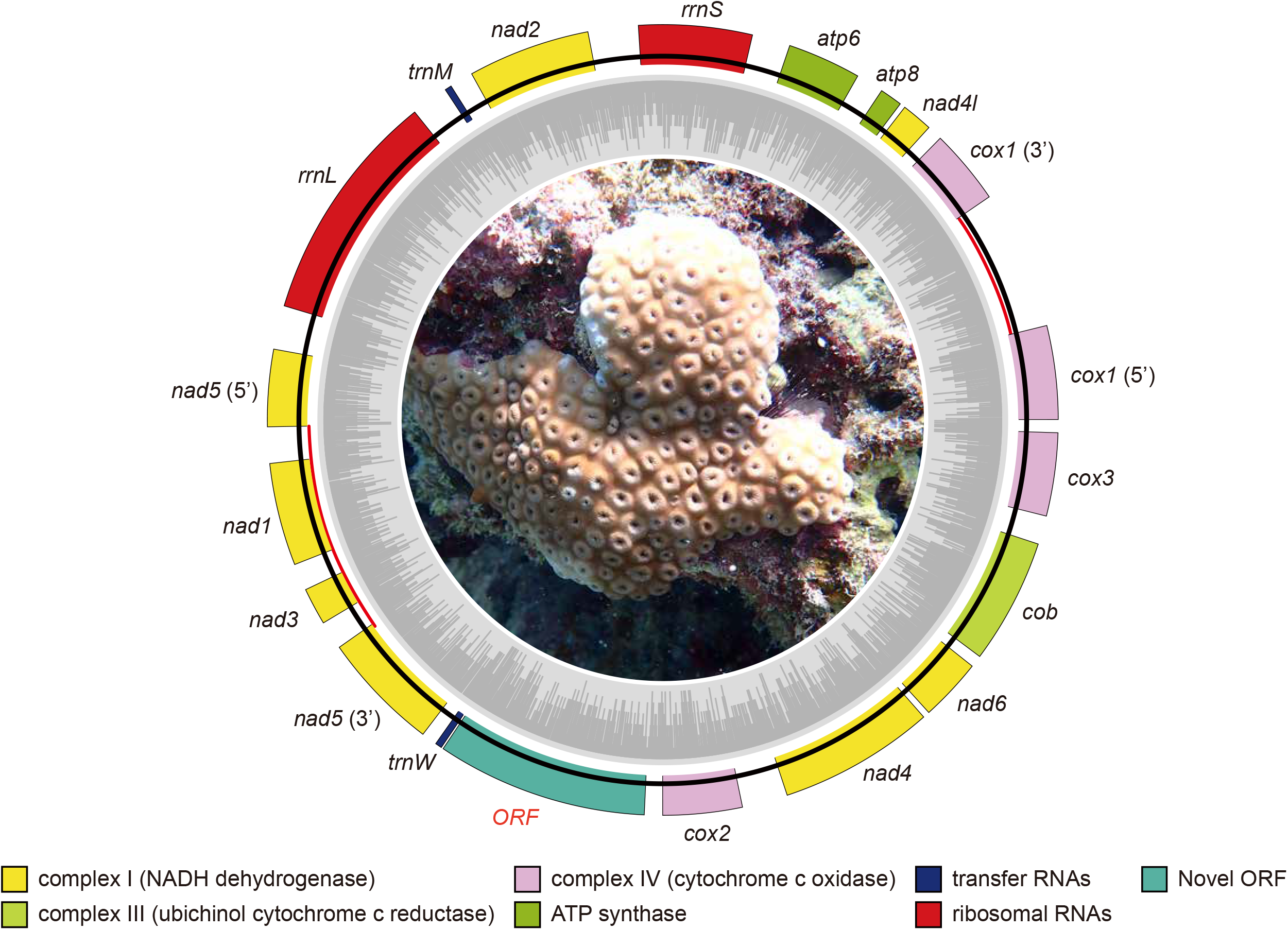
The complete mitochondria genome of *Palythoa tuberculosa*. Inner circles (grey) indicate GC contents (%). Inner red line indicates intron for *nad5* and *cox1*. NADH dehydrogenase (yellow), ubichinol cytochrome c reductase (light green), cytochrome c oxidase (pink), ATP synthase (green), tRNAs (blue), rRNAs (red), and novel open reading frame (mint green). Photo was taken by Megumi Kanai.

Codon usage was not even across the PCGs, with codons for arginine, leucine, or serine occurring more frequently than those for other amino acids (Figure 2). Such biased synonymous codon usage has also been reported in other cnidarians (e.g., scleractinian corals; Baeza and Rosales, 2025). The Ka/Ka ratios for all mitochondrial PCGs in *P. teberculosa* exhibited values were less than 1, or most cases showed no substitution in pairwise comparisons with *P. mutsuki* (Supplementary Table S1), indicating that these genes are evolving under purifying selection.

**Figure 2.**
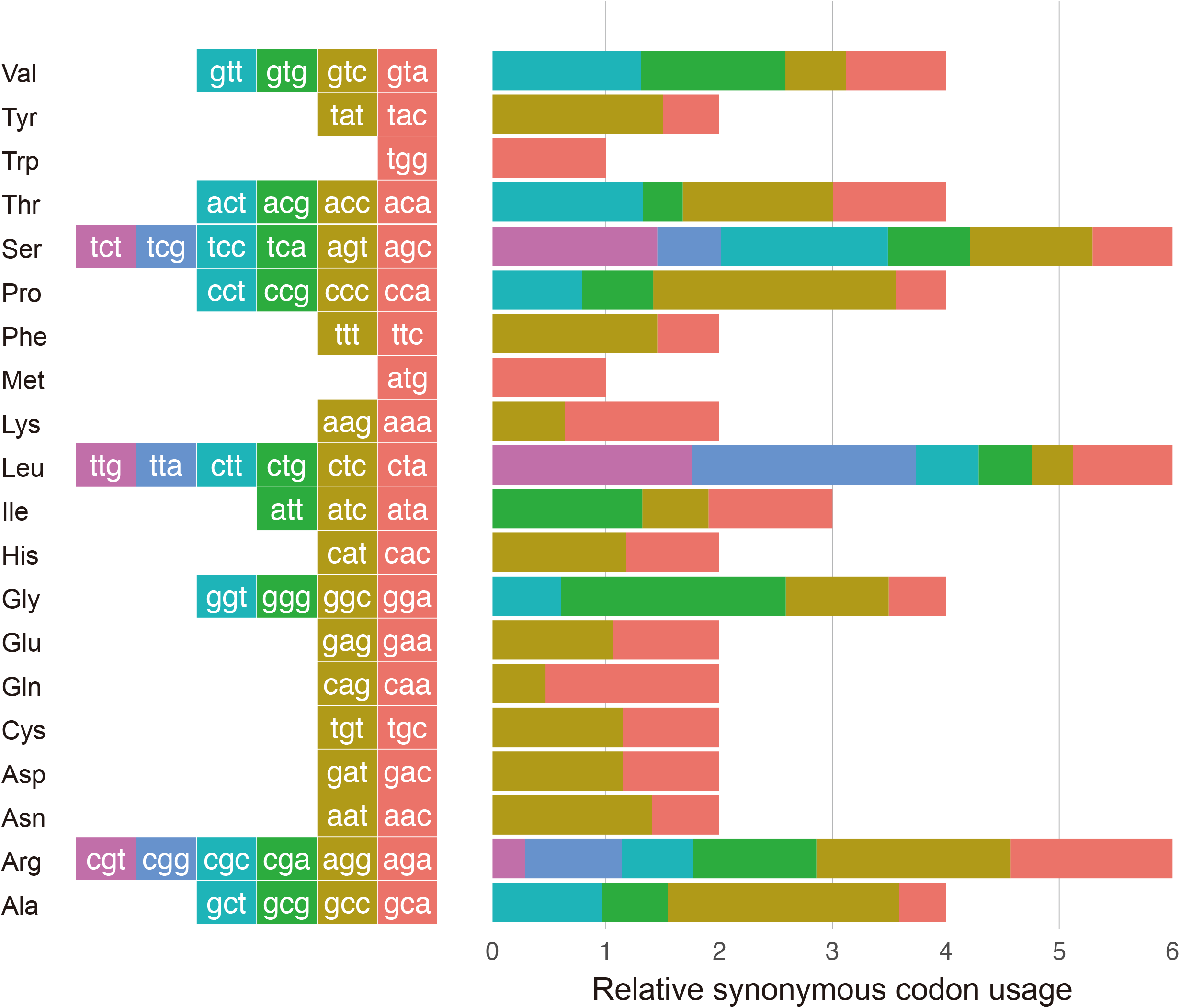
Graphical depiction of codon usage in the protein-coding genes of *Palythoa tuberculosa* mt-genome. Relative abundances of codons are associated by color within each bar of their respective amino acids. *atp6, atp8, cob, cox1*–*3, nad1*–*6*, and *nad4l* were used in this analysis.

A maximum-likelihood phylogenetic tree was constructed (Figure 3), and tree topology was identical to that previously reported by Fourreau et al. (2023), confirming the robustness of the phylogenetic relationships. Although extensive mitochondrial gene order rearrangements are often observed in cnidarians, no changes were detected within *Palythoa* and *Zoanthus* (Figure 3), in agreement with earlier studies (Chi and Johansen, 2017; Fourreau et al., 2023; Sinniger et al., 2007). Synteny analysis further revealed that the novel ORF between *trnW* and *cox2* accounted for the increased mt-genome size in *P. tuberculosa* (Figure 4). In addition, this insertion was also present in *P. caribaeorum* with complete sequence identity and identical gene order (Figure 4; Supplementary Figure S1), suggesting an origin in their common ancestor.

**Figure 3.**
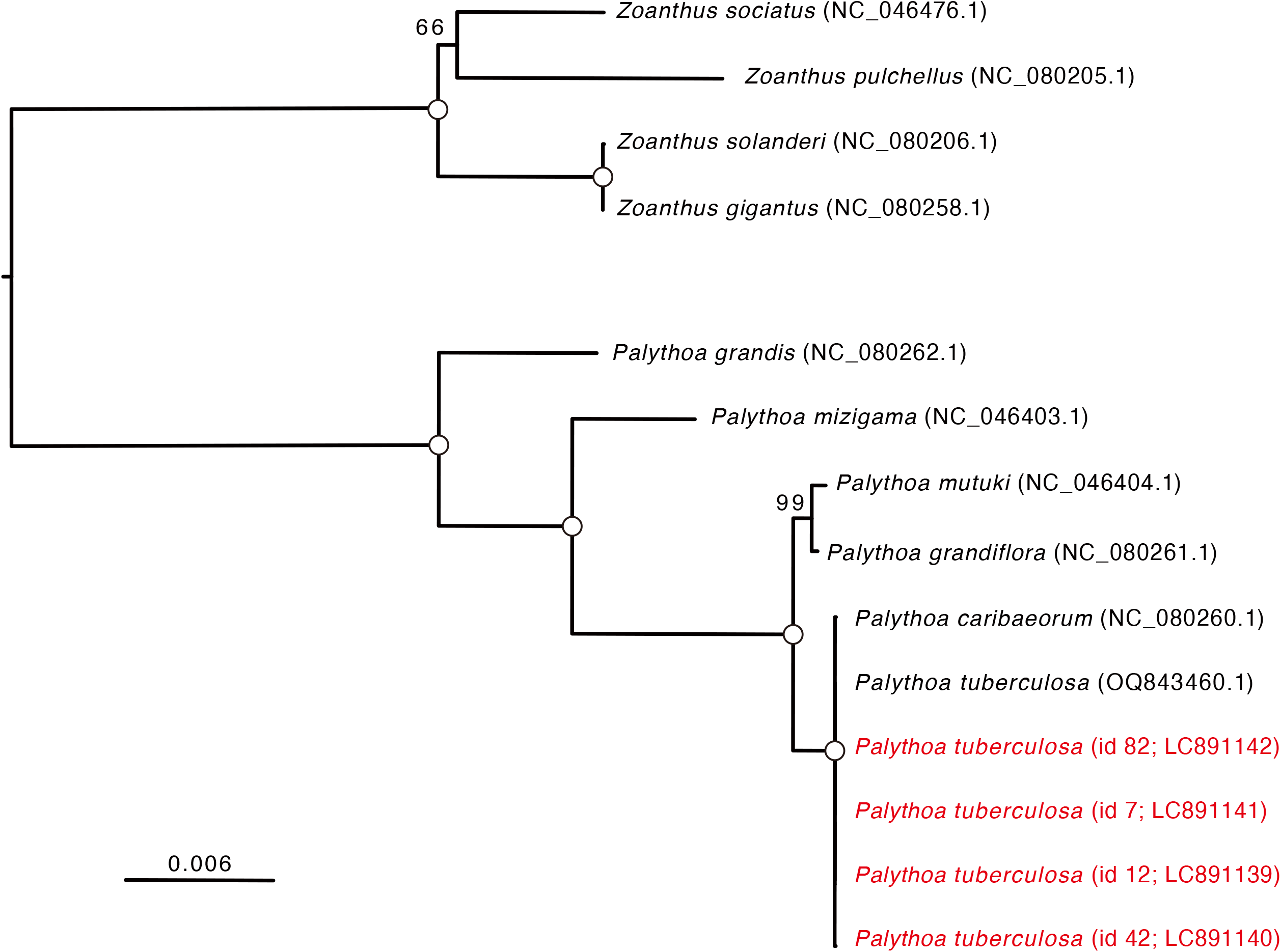
A molecular phylogenetic tree based on mitochondrial genomes. Tree topology was estimated based on 25,302 nucleotide positions. Species belonging to the genus *Zoanthus* (Family: Zoanthidae) were used as outgroup. Accession numbers for mitochondrial genomes are shown in parentheses after scientific names. The bar indicates expected substitutions per site in aligned regions. Open circles indicate 100% bootstrap support (1,000 replicates).

**Figure 4.**
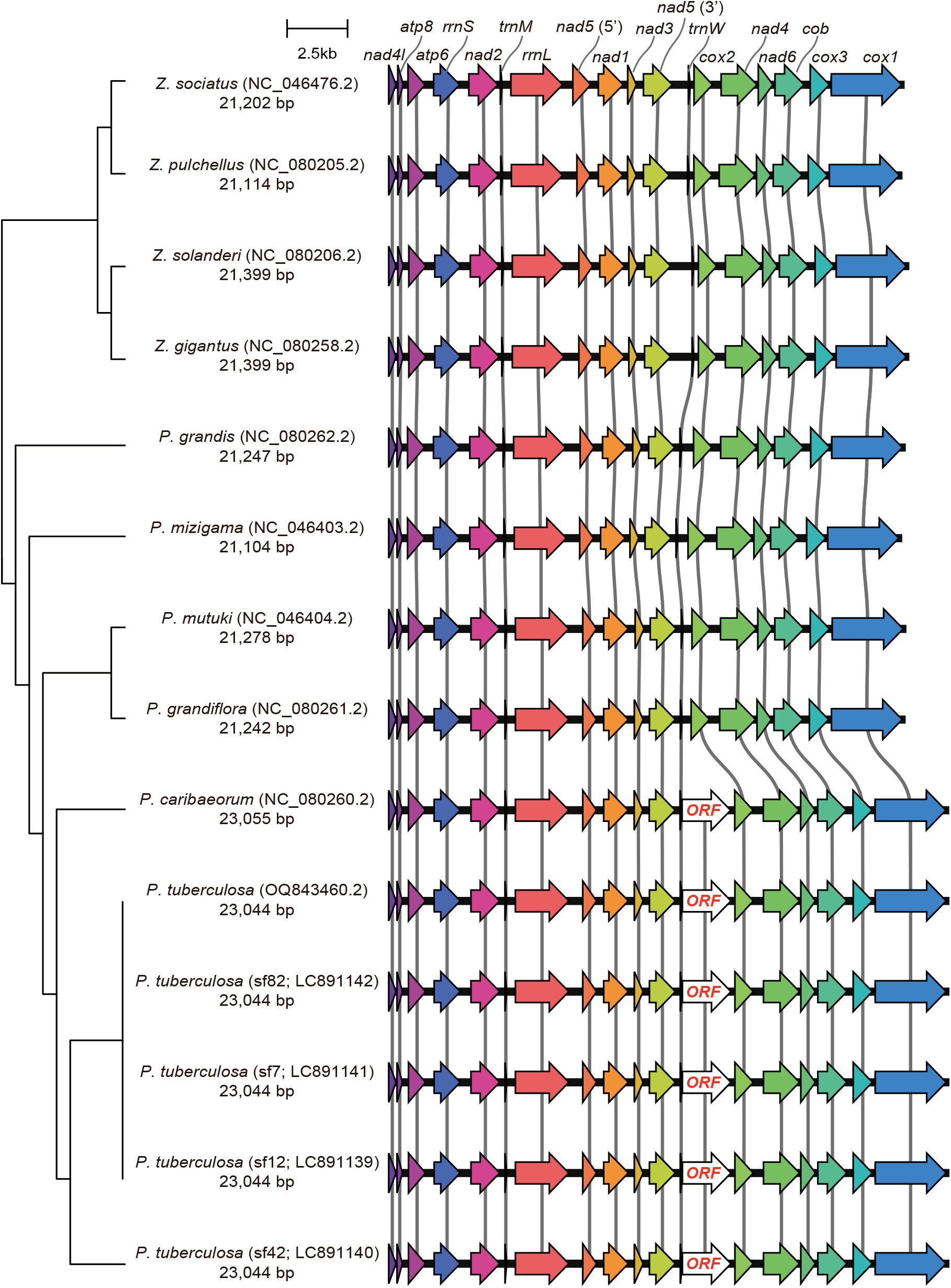
Mt-genome synteny among *Palythoa* species. Tree topology shows phylogenetic relationship inferred in Figure 1. Each gene is colored and orthologs are connected by lines. “ORF” indicates open reading frame found in this study. Arrowheads indicate transcription directions. The bar indicates 2.5 Kbp. Accession numbers and Mitochondrial genome sizes are shown along with scientific names.

To investigate the evolutionary origin of this ORF, we performed sequence homology search against NCBI clustered NR database (accessed on 2025-09-25). Although partial blast hit to mitochondrial genes of sea anemones was detected (Supplementary Table S2), the low sequence identity (∼30%) suggests a lineage-specific origin during evolution. ORF extensions (up to 146 amino acids) have been reported in several mitochondrial genes of the sister genus *Zoanthus*, relative to scleractinian corals and sea anemones (Chi et al., 2017). The novel ORF insertion identified in this study represents the first case in zoantharians, distinct from previously reported instances of ORF extension. A comparable instance of novel ORF insertion has been described in the coral genus *Pocillopora* (Order Scleractinia: Family Pocilloporidae) (Flot and Tillier, 2007), although its evolutionary origin remains unresolved. In *Pocillopora*, ORF accumulates nucleotide substitutions and serves as a marker for species identification (Nakajima et al., 2017), implying that ORFs in mitochondrial genomes may play important lineage/species-specific roles.

One mechanism proposed for mitogenome expansion is the tandem duplication–random loss (TDRL) model (Moritz et al., 1987), which posits that duplication of a gene block—arising through replication errors such as strand slippage, mispairing, or mis-initiated replication—is followed by random deletion of redundant copies. This mechanism typically preserves similarity between duplicated and original genes; however, the novel ORF identified here shares no detectable similarity with other mitochondrial genes (Supplementary Table S3), suggesting the possibility of *de novo* gene emergence or accelerated substitution accumulation. Further transcriptomic and experimental analyses will be required to elucidate the functional significance of this ORF in *P. tuberculosa* and *P. caribaeorum*.

## Conclusions

This study reported the complete mitochondrial genomes of *P. tuberculosa* with comprehensive gene annotation. We identified a novel ORF insertion to the mt-genome of this species. Phylogenetic analysis corroborated established evolutionary relationships within the group, while synteny analysis revealed that the novel ORF insertion contributed to the enlarged genome size. Although the function of this ORF remains unknown, future studies incorporating broader taxon sampling across zoantharians, together with functional investigations, will provide deeper insights into the evolutionary dynamics of mitochondrial genome evolution in cnidarians.

## Acknowledgements

We thank members of the Sequencing Section of OIST for conducting genome sequencing and members of the Scientific Computing and Data Analysis section of OIST for computing resources. We also thank Mr. Shogo Gishitomi and Ms. Natsuki Watanabe of Incorporated Foundation Okinawa Environment and Science Centre for conducting DNA extraction.

