## Supplementary Figures for "The mitochondrial genome of *Palythoa tuberculosa* (Esper, 1805) contains a novel ORF with unknown function"

Corresponding author:

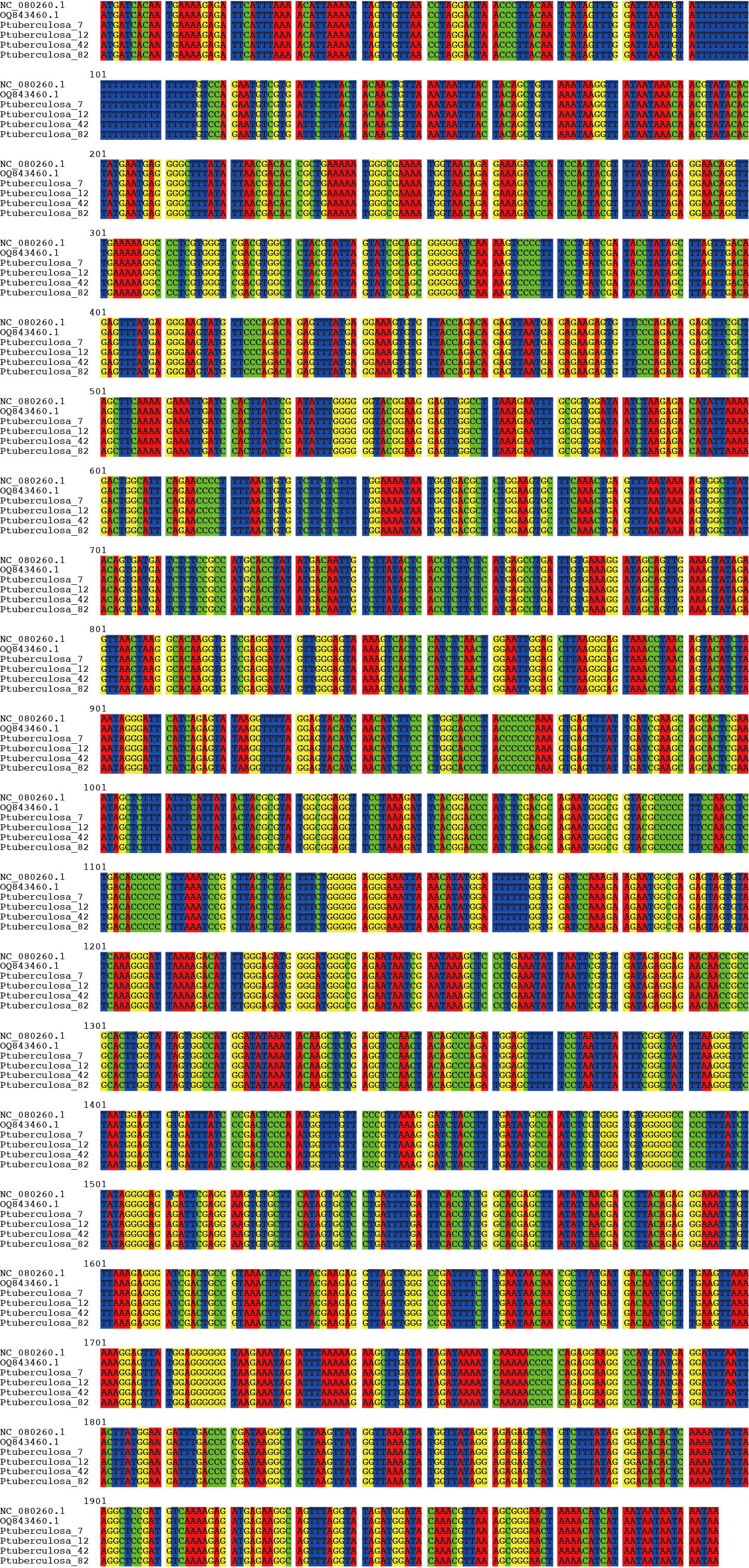

**Supplementary Figure S1.** **Alignment of the novel open reading frame in mt-genomes of *Palythoa tuberculosa* and *Palythoa caribaeorum*.**

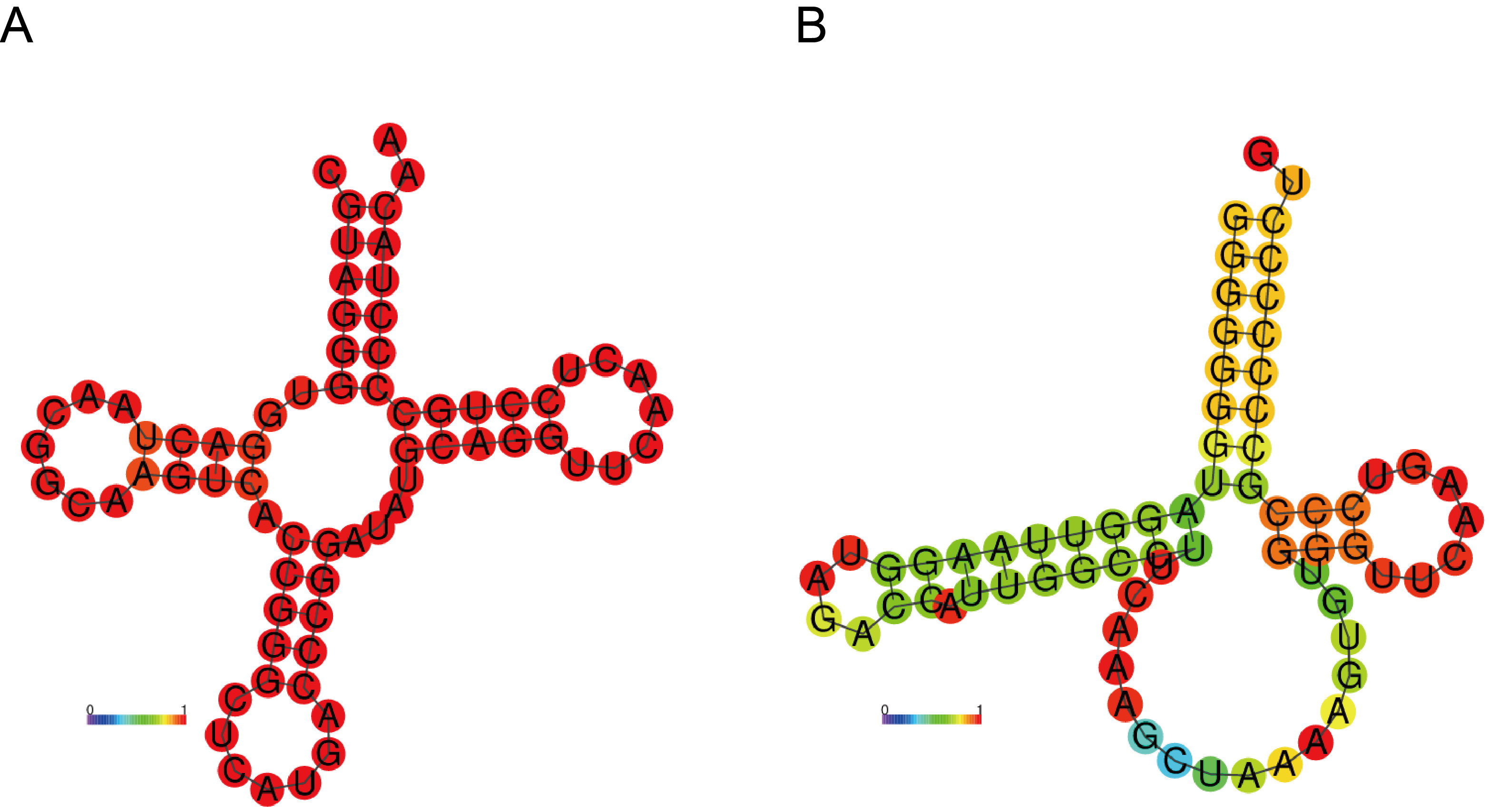

**Supplementary Figure S2. Secondary structure of *trnM* (A) and *trnW* (B).**

Color represents base-pair probabilities calculated with RNAfold web server.

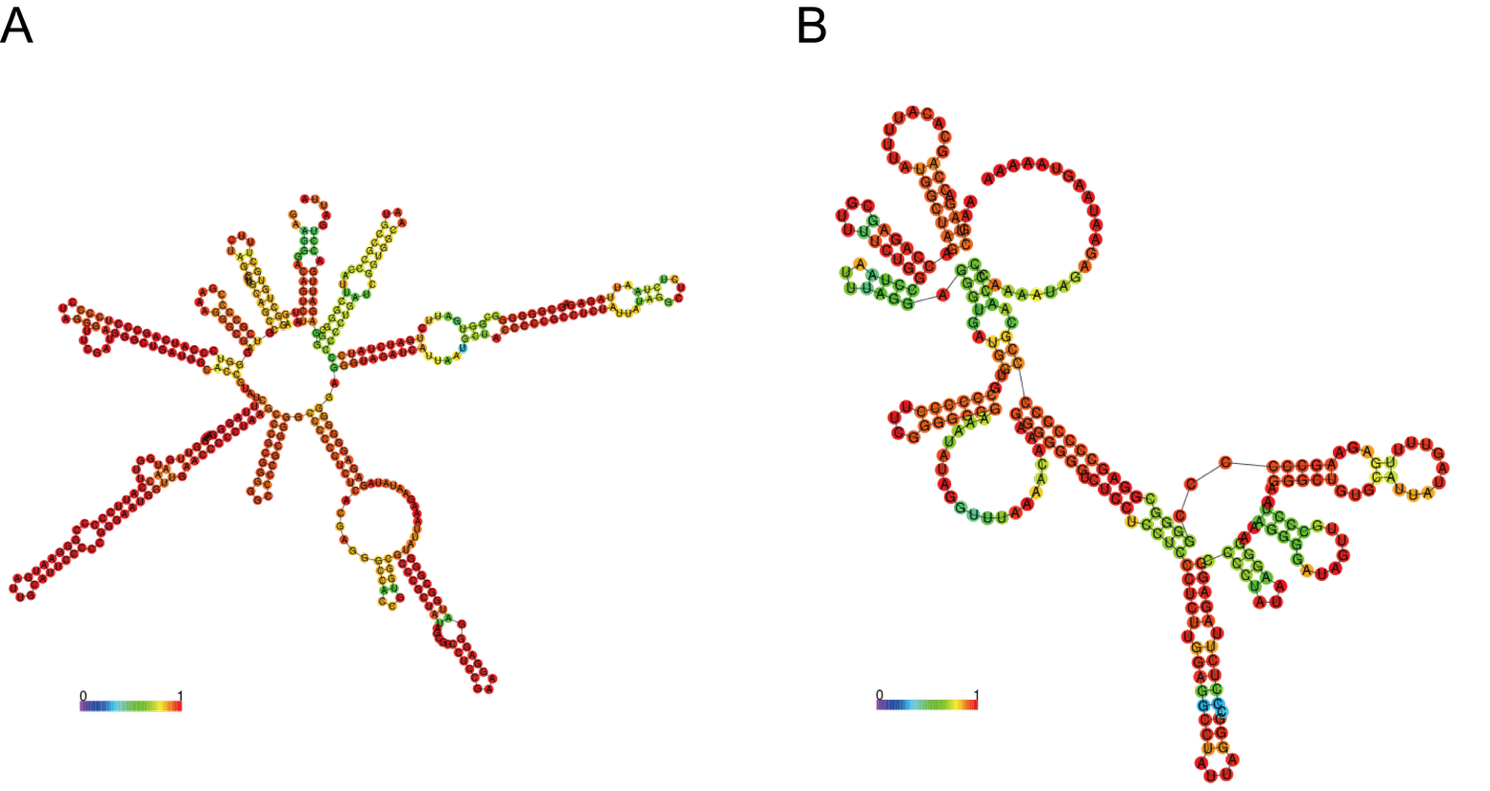

**Supplementary Figure S3. Secondary structure of putative control regions located between *cox2* and *nad4* (A) and between *cob* and *cox3* (B).**

Color represents base-pair probabilities calculated with RNAfold web server.
